# A primitive type of renin-expressing lymphocyte protects the organism against infections

**DOI:** 10.1101/770511

**Authors:** Brian C. Belyea, Araceli E. Santiago, Wilson A. Vasconez, Vidya K. Nagalakshmi, Theodore C. Mehalic, Maria Luisa S. Sequeira-Lopez, R. Ariel Gomez

## Abstract

The hormone renin plays a crucial role in the regulation of blood pressure and fluid-electrolyte homeostasis. Normally, renin is synthesized by juxtaglomerular (JG) cells, a specialized group of myoepithelial cells located near the entrance to the kidney glomeruli. In response to low blood pressure and/or a decrease in extracellular fluid volume (as it occurs during dehydration, hypotension, or septic shock) JG cells respond by releasing renin to the circulation to reestablish homeostasis. Interestingly, renin-expressing cells also exist outside of the kidney, where their function has remained a mystery. We discovered a unique type of renin-expressing B-1 lymphocytes that may have unrecognized roles in defending the organism against infections. These cells synthesize and release renin, entrap and phagocyte bacteria and control bacterial growth. The ability of renin-bearing lymphocytes to control infections – which is enhanced by the presence of renin – adds a novel, previously unsuspected dimension to the defense role of renin-expressing cells, linking the endocrine control of circulatory homeostasis with the immune control of infections to ensure survival.

## INTRODUCTION

Renin-expressing cells emerged in nature over 400 million years ago(1, 2). Throughout evolution, they have acquired numerous defensive functions that rendered them as perfect machines to ensure our survival in response to a variety of homeostatic threats(3). They control blood pressure, fluid-electrolyte balance, vascular development, glomerular regeneration and may participate in the regulation of oxygen delivery to tissues(1, 3). Although renin cells were first discovered in the juxtaglomerular areas of the adult kidney arterioles, from where they release renin to the circulation to regulate blood pressure and fluid-electrolyte homeostasis, their appearance in this organ is a late event in their developmental history(1). In fact, during early mouse and human development, cells that express renin emerge in multiple tissues and organs before they appear in the kidney(1). The function of renin cells outside the kidney has remained a mystery and the subject of great speculation. We report here the discovery of a primitive type of renin-expressing cell within hematopoietic organs that persists throughout adulthood and may have hitherto unrecognized roles in defending the organism against infections. These cells possess unique capabilities to trap and phagocyte bacteria and control bacterial growth. The ability of renin-bearing lymphocytes to control infections adds a novel and unsuspected dimension to the defense role of renin-expressing cells.

## RESULTS AND CONCLUSIONS

Using lineage-tracing, flow cytometry, fluorescence imaging, and gene expression analysis, we examined the temporal appearance, distribution, identity, and evolution of renin progenitors throughout hematopoietic ontogeny. Renin-expressing, hematopoietic progenitors first appear within the yolk sac during mid-gestation. Using reporter mice (*Ren1*^*dcre*/+^; *mTmG*), which express GFP in all cells that expressed renin and their descendants(4–6), we identified GFP^+^ / renin^+^ cells first within the yolk sac at E11.5 (Figs. 1a, b). Their presence within this organ is transient and not found beyond E13.5. These dual hematopoietic and renin-bearing precursors leave the yolk sac and colonize the fetal liver and spleen at ~E15.5 (Fig. 1c). As embryonic development progresses, the cells increase in number within the fetal liver and spleen, representing approximately 10% of hematopoietic cells within these tissues at the time of birth (Figure 1d, e). To define the identity of these cells, we immunophenotyped them using flow cytometry and a panel of well-characterized cell surface antibodies(7–9). In the fetal liver and spleen, the renin lineage cells express B lymphocyte cell surface markers CD19 and CD43, however they have dim B220 expression and are negative for a cocktail of lineage markers (“Lin”), consistent with a B-1 progenitor immunophenotype (B220^dim^CD19^+^CD43^+^Lin^−^) (10) (Fig. 1f).

**Figure 1.**
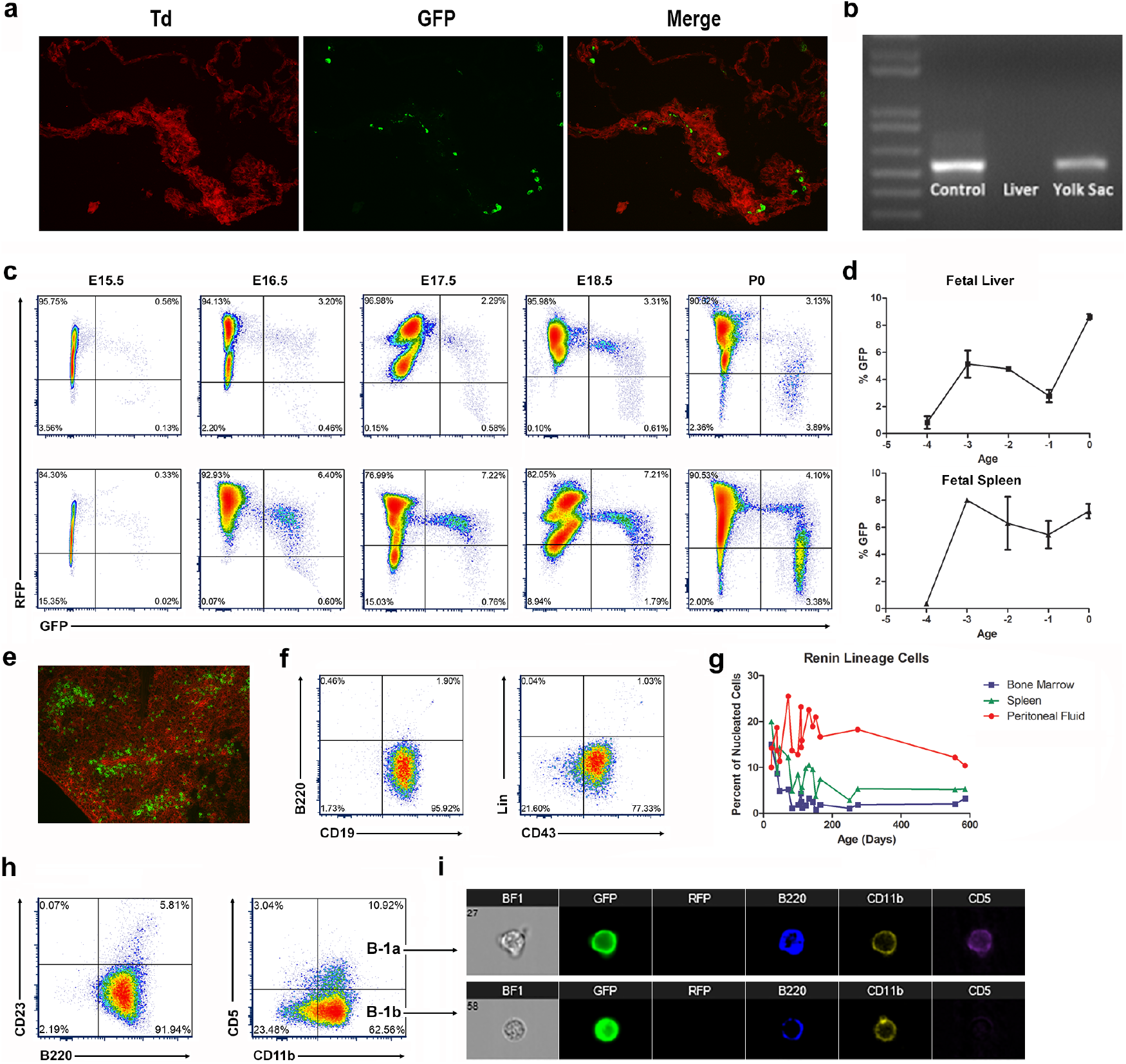
Renin-expressing cells arise during mid-gestation as B-1 progenitor cells and persist throughout adult life in the peritoneal cavity as B-1 B cells. a. Renin progenitors appear in the yolk sac at E11.5 as individual GFP^+^ cells within the Td^+^ yolk sac tissue of *Ren1*^*dcre*/+^;*mTmG* mice. b. Semi-quantitative RT-PCR confirmed renin expression in the yolk sac at E11.5. Newborn kidneys and E11.5 livers were used as positive and negative controls respectively. c. Flow cytometry representative plots show that renin-lineage cells (GFP^+^, x-axis) appear in the fetal liver and spleen at E15.5. With advancing gestational age (E15.5 – P0, left to right), there is an increase in the percentage of GFP^+^ cells in both the fetal liver (top panel) and fetal spleen (bottom panel). d. Percentage of renin progenitors from E15.5 (−4) to P0 using flow cytometry. e. Renin lineage (GFP^+^) cells are present in the spleen within the white pulp in newborn mice. f. Renin-expressing cells in the fetal tissues show a B-1 progenitor phenotype (CD19^+^, B220^dim^, CD43^+^, and Lin^−^). g. The percentage of renin-lineage cells decreases in the bone marrow and spleen over time but persists in the peritoneal cavity throughout adult life. h. Renin lineage (GFP^+^) cells in the peritoneal cavity are B-1b B cells (B220^+^, CD23^−^, CD11b^+^, CD5^−^) and B-1a B cells (B220^+^, CD23^−^, CD11b^+^, CD5^+^). i. IMAGESTREAM flow cytometry shows that renin-lineage (GFP^+^) peritoneal cells express B220, CD11b, and +/− CD5. Cells that express CD5 are B-1a B cells, and cells that do not express CD5 are B-1b B cells.

Following birth, renin lineage cells are found throughout the hematopoietic system, including bone marrow, spleen, and peripheral blood. Because B-1 cells have been observed in serosal compartments(7),(11) we confirmed that the mouse peritoneal cavity harbored renin-expressing cells with a lymphocyte pedigree (Fig. 1g). Whereas the proportion of renin cells in the peritoneal cavity is maintained throughout adult life, their proportion in the bone marrow and spleen diminishes with age (Fig. 1g). Renin lineage (GFP^+^) cells from the bone marrow, spleen, and peripheral blood are B-2 B lymphocytes (B220^+^CD19^+^CD23^+/−^CD11b^−^)(4). However, renin progenitors in the peritoneal cavity are B-1 B cells (B220^dim^CD23^−^CD11b^+^CD5^+/−^) (Fig. 1h, i).

To confirm that renin-expressing B-1 progenitors in the fetal hematopoietic tissues give rise to mature B-1 cells in the adult, we performed transplant studies. We isolated hematopoietic cells from the fetal livers of E16.5 *Ren1*^*dcre*/+^;*mTmG* reporter mice and injected these cells into the tail vein of irradiated wildtype adult mice. Transplant of fetal liver cells gave rise to GFP^+^ B cells within the bone marrow, spleen, peripheral blood, and peritoneal cavity of recipient mice. Indeed, the percentage of GFP^+^ cells in the bone marrow and spleen of transplant recipient mice was similar to adult reporter mice (Supplemental Fig. 1a,b). The phenotype of the renin-lineage cells within the transplant recipients was also similar to that of the age-matched reporter mice (Supplemental Fig. 1a,b). Thus, these transplant studies confirm that renin-expressing hematopoietic progenitors during fetal life give rise to B-1 and B-2 cells in adult animals.

Until recently, the origin of B1 lymphocytes has been unclear(12). Our studies indicate that B1 lymphocytes which had expressed renin during early development, originate within the yolk sac during the initial wave of primitive B lymphopoiesis and then expand to the fetal liver and spleen prior to the development of definitive hematopoiesis. Those cells persist during adult life as B-1 B cells in the peritoneal cavity and, to a lesser extent, as B-2 B cells in the bone marrow, spleen, and peripheral blood.

To determine whether these lymphocytes with a renin pedigree manufacture and release renin, we used a dot membrane immunoassay(13, 14). Peritoneal cells from *Ren1*^*dcre*/+^; *mTmG* reporter mice were cultured on nitrocellulose membranes for 48 hours followed by dot immunoassay for renin using a well characterized renin antibody and color development detection kit as previously described(15). Figure 2a shows that peritoneal cells derived from the renin lineage appear as discrete blue dots indicating that these cells actively release renin. A similar pattern is obtained with As4.1 cells, mouse tumoral cells that secrete renin constitutively (Fig. 2a). No spots were detected in C2C12 cells, skeletal muscle cells that do not normally manufacture renin (Fig. 2a). Further, peritoneal cells from WT mice expressed authentic renin mRNA, verified by sequencing, and not a related aspartyl protease (Fig. 2b). Renin mRNA was not present in cells from renin null mice(16). To further document these findings, we obtained peritoneal cells from *Ren1*^*c-YFP*^ mice(17), where YFP is under the control of the renin super-enhancer(3, 18) and reports the transcriptional activity of the renin gene. Figure 2c shows YFP+ cells from the peritoneal cavity, indicating that these cells actively transcribe renin and YFP.

**Figure 2.**
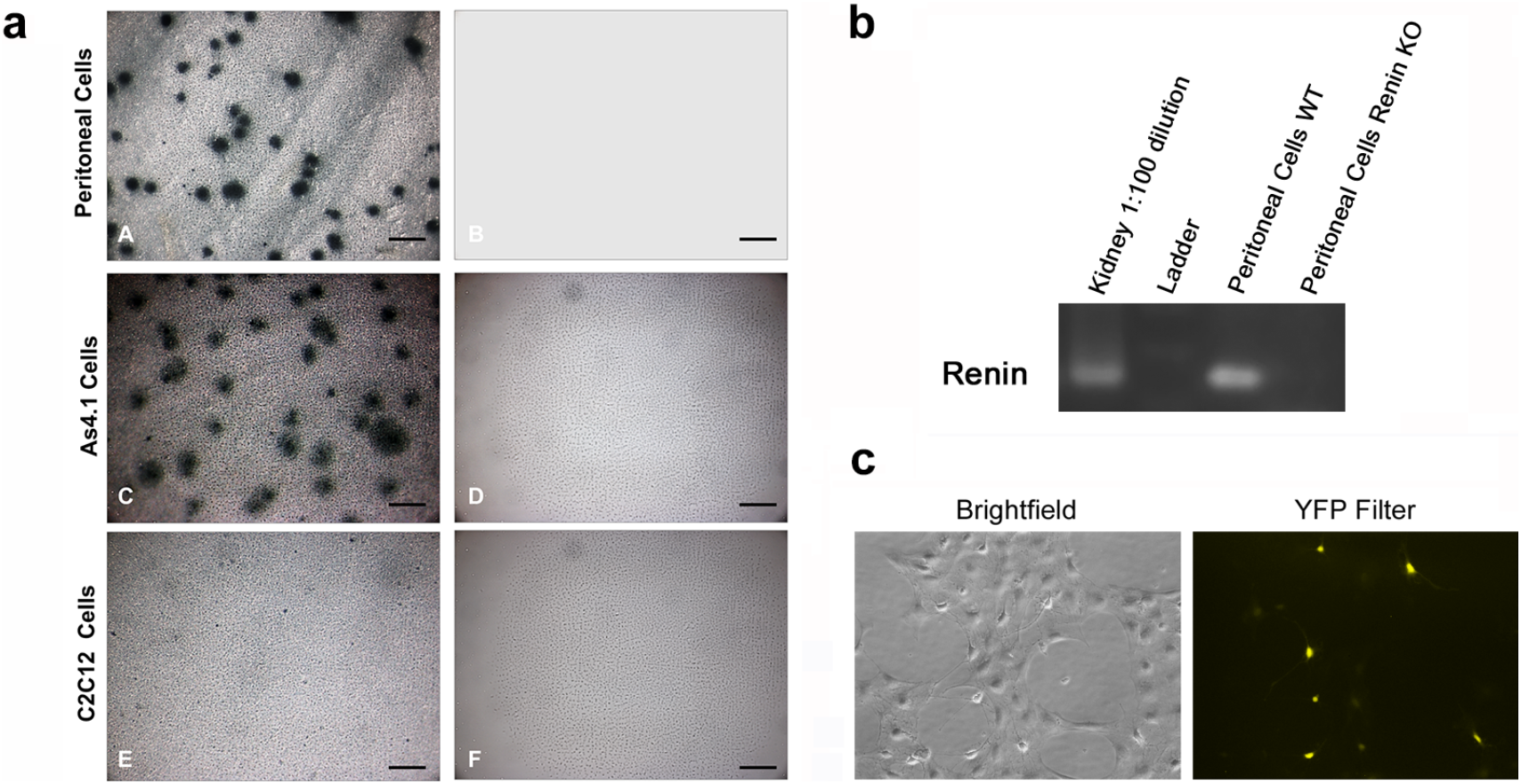
Peritoneal Cells Express Renin. a. A dot membrane immunoassay shows that peritoneal cells derived from the renin lineage appear as dark blue dots, indicating that these cells actively manufacture and release renin (A). A similar pattern is obtained with As4.1 cells, a mouse tumoral cell line that secretes renin constitutively (C). By the contrary, no spots were detected in C2C12 cells, skeletal muscle cells that do not normally synthesize renin (E). Further, no spots are detected in any of the cells when the membrane immunoassay is performed in the absence of the primary renin antibody (B, D, F) (scale bars: A-F 200 μm). b. Semi-quantitative RT-PCR was performed on wildtype peritoneal cells and peritoneal cells from a renin KO animal. Kidney RNA was used as a positive control. c. Peritoneal cells from Ren1^c-YFP^ reporter mice, where YFP marks active renin expression were grown in culture. YFP was demonstrated by immunofluorescence.

To examine whether renin-bearing GFP^+^ B1 lymphocytes interact with bacteria, we incubated them with CFP-expressing *E. Coli.* GFP^+^ B-1 lymphocytes trailed the CFP^+^ bacteria and made numerous contacts with them via the assembly and disassembly of pseudopod-like extensions (See extended data, video 1). To explore the nature of the lymphocyte-bacterial contacts, we used scanning electron microscopy. Peritoneal cells emit cable-like extensions and mesh-like structures that entrapped the bacteria which subsequently become immobilized (Fig. 3a). To our knowledge, these poorly understood structures have not been previously described in lymphocytes.

**Figure 3.**
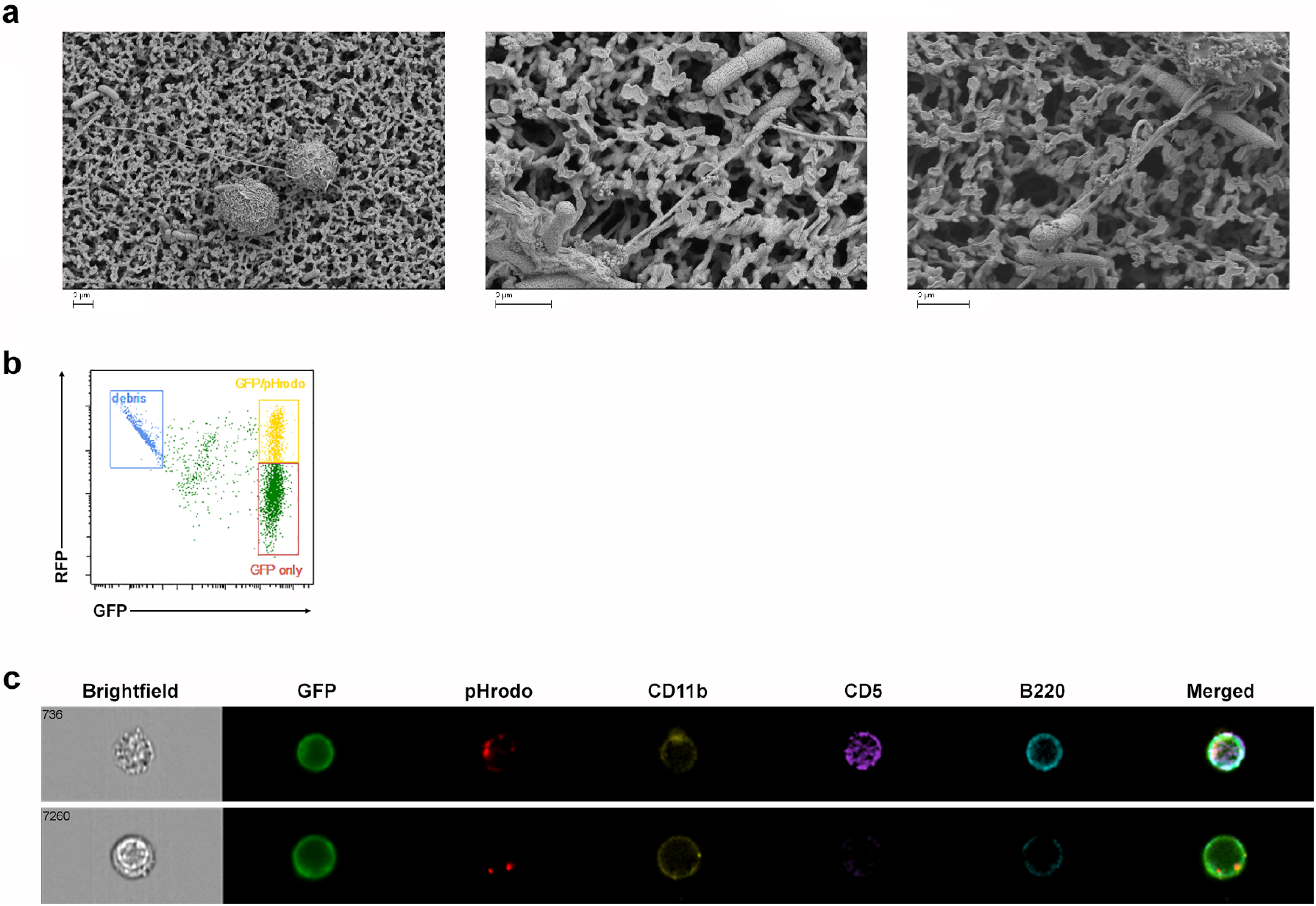
Renin-lineage peritoneal cell physically interact with bacteria and phagocytose bacteria particles. a. Renin-lineage cells were incubated with 042 E. Coli bacteria for 2 hours and then processed for scanning electron microscopy. Renin-lineage B-1 cells associate with bacteria through cable-like extensions. b. A representative flow cytometry plot demonstrating GFP^+^ cells (renin lineage cells) which are double positive for pHrodo. c. Identification of renin lineage peritoneal cells that phagocytosed *E. Coli* particles using an ImageStreamX Mark II system. The ImageStream system software (IDEAS) obtained images of events determined to be GFP^+^ cells that have phagocytosed pHrodo. The top panel shows an image of a renin-lineage B-1a B cell which has phagocytosed *E. Coli* particles (pHrodo^+^). This cell expressed CD11b, CD5, and B220. The bottom panel shows an image of a renin-lineage B-1b cell (CD5-).

To determine whether renin-lineage B-1 cells phagocyte bacteria, we exposed GFP^+^ B-1 cells to *E. Coli* particles labeled with pHrodo(19). When the particles are internalized and routed to lysosomes, the low pH in the lysosomes results in red fluorescence emanating from the bacterial particles. Using ImageStream software (IDEAS), we observed those phagocytic events. Figure 3b shows the dot-plot of intensity of GFP (X axis) versus pHrodo (Y axis). There is a double-positive GFP/pHrodo population indicating that a significant proportion of GFP^+^ B-1 cells have ingested the bacterial particles (Fig. 3b, yellow box). Figure 3c shows examples of renin lineage B1a (top) and B1b (bottom) lymphocytes which have ingested bacterial particles.

To determine whether renin-expressing B-1 lymphocytes possess bactericidal activities, we incubated them with *Salmonella typhimurium* (ATCC14028) for two hours and measured bacterial growth over 24 hours by bacterial counting of colony forming units. Incubation of bacteria with B-1 cells markedly diminished the number of colonies compared to bacteria growing in the absence of B-1 cells, indicating that B-1 cells possess the ability to inhibit bacterial growth (Fig. 4a). To explore whether renin produced by B-1 cells contributed to the antimicrobial function, we cultured peritoneal B-1 lymphocytes from renin null (*Ren1*^*c−/−*^) mice and from wild-type controls with *Salmonella* at various cell : bacteria ratios for two hours. Bacteria co-cultured with wildtype B-1 cells had decreased growth. However, when bacteria were co-cultured with renin null B-1 cells, the inhibition of bacterial growth was markedly diminished (Fig. 4a, b) suggesting that intracellular renin may play a role in defending against bacterial pathogens. Together, these studies uncovered an unsuspected, previously unrecognized role of renin-expressing cells in the defense of the organism against infections.

**Figure 4.**
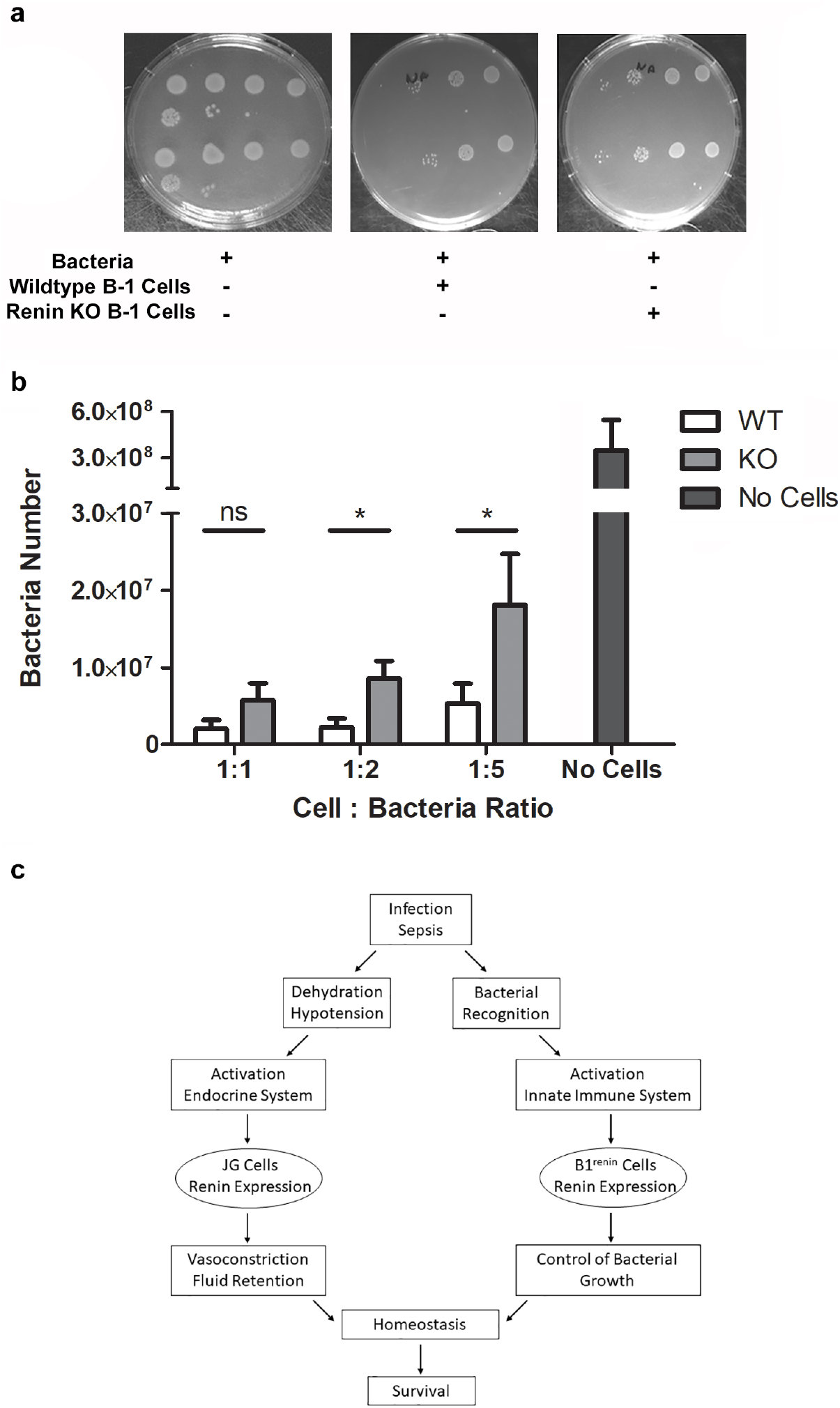
Peritoneal cells from renin KO mice have decreased ability to inhibit bacterial growth in vitro. a. Peritoneal B-1 lymphocytes were obtained from wildtype and renin-KO mice. These cells were co-cultured with *Salmonella typhimurium* bacteria for 2 hours. A third group included *Salmonella typhimurium* bacteria alone. Cells were removed after two hours, and the bacteria were grown in agar for 24 hours. Bacteria numbers were then calculated. b. These experiments were repeated at cell : bacteria ratios of 1:1, 1:2, and 1:5. Each experiment was performed 5 times, and the results were grouped together. c. Schematic as to how the endocrine renin-angiotensin system and the innate immune system coordinate efforts in response to threats to homeostasis.

We have previously shown that progenitor cells from different embryonic layers express renin during development(1). Those progenitors differentiate into a variety of phenotypically and functionally diverse group of cells distributed throughout the body. Whereas most of them stop producing renin, the kidney juxtaglomerular cells and the B-1 lymphocytes retain the ability to synthesize and release renin in adult life(1). Interestingly, these two cell types share some core transcriptional regulators(4). We have previously shown that deletion of RBP-J in the mouse kidney alters the fate of renin cells. As a result, the animals are unable to maintain blood pressure when exposed to a threat to homeostasis(20). Similarly, deletion of RBP-J in pre-B lymphocytes leads to inability of the cells to differentiate resulting in their uncontrolled neoplastic proliferation(4). Kidney juxtaglomerular cells and B1 lymphocytes may also share functional responses. For instance, administration of captopril which decreases blood pressure in adult mice, results in an increase in the number of renin producing cells both in the kidney and in the bone marrow(4) suggesting that these two seemingly distant cells act in concert to maintain homeostasis when confronted with a threat to survival. A further look at these cells indicate them to be engaged in the unique specialized control of homeostatic defense. JG cells in the kidney located at the vascular pole of glomeruli are strategically located to sense and respond to changes in the composition and volume of the extracellular fluid and to changes in blood pressure. Similarly, B1 lymphocytes recognize and react to the presence of foreign antigens and microorganisms. Figure 4C illustrates how these cells may coordinate their response upon a threat to survival, as it frequently occurs in gastrointestinal or abdominal infections. Under these circumstances, hypotension and volume depletion induces JG cells to release renin to the circulation leading to the generation of angiotensin(s), vasoconstriction, sodium chloride reabsorption and reestablishment of fluid electrolyte and blood pressure balance(21). Similarly, peritoneal B-1 renin cells which have the capability to recognize, entrap, phagocyte and kill bacteria, act as an early line of defense to counteract and stop the threat. Interestingly, the presence of renin renders B-1 lymphocytes more effective in bacterial killing, suggesting that renin per se may have an added role in the degradation of bacterial components.

Thus, renin, and the diverse group of cells that synthesize it, are at the epicenter of two systems crucial for survival: the endocrine renin-angiotensin system driven to maintain cardio-circulatory homeostasis by regulating volume and tissue perfusion and the innate immune system designed for the rapid control of infections.

## METHODS (Complete Methods included in Supplemental Data)

### Mice

*Ren1*^*dCre*/+^ mice express Cre recombinase in renin cells(6). *Ren1*^*dcre*/+^ mice were bred to the lineage reporter line *mT/mG*, in which non-recombined cells express mTd and Cre-mediated recombined cells express GFP(22). Thus, GFP is expressed in renin-expressing cells and all descendants. *Ren1*^*c-YFP*^ transgenic mice express YFP in cells that actively express renin and were used to identify cells that actively express renin(17). *Ren1*^*c−/−*^*Ren1*^*c-YFP*^ mice are renin knockout mice which also express YFP in cells that attempt to express renin.

### Cells

Peritoneal cells were obtained from WT (*Ren1*^*c-YFP*^) and renin knockout (*Ren1*^*c−/−*^*Ren1*^*c-YFP*^) mice. Briefly, mice were anesthetized, and scissors and forceps were used to remove the skin overlying the ventral portion of the peritoneum. 5 mL of PBS with 3% FBS was injected into the peritoneal cavity with a 30 gauge needle. The peritoneum was then massaged for 5 minutes to mobilize cells into the PBS solution. An 18 gauge needle was then used to remove the fluid containing peritoneal cells. The cell suspension was then pelleted and resuspended in DMEM + 5% FBS media. As4.1 cells are a renin-expressing tumoral cell line (ATCC, CRL-2193)(23). C2C12 cells are an immortalized mouse myoblast cell line (ATCC, CRL-1772).

### Bacteria

DH5α E. coli bacteria were co-cultured with peritoneal cells from *Ren1*^*dCre*/+^;*mTmG* mice. DH5α bacteria were transformed with a CFP plasmid (Oxford Genetics Ltd, Oxfordshire, United Kingdom) using Kanamycin selection (1μl of CFP plasmid added to 50 μl bacteria). Colonies positive for CFP were selected. These bacteria were subsequently termed “DH5α-CFP”. DH5α-CFP bacteria were co-incubated with peritoneal cells, and time-lapsed pictures were taken. 042 bacteria are a strain of enteroaggregative *Escherichia coli(24)*. These bacteria were used for SEM (below). For bacterial ingestion and killing assays, we used *Salmonella typhimurium* (ATCC 14028) and modified the protocol as described(25, 26). Briefly, we isolated peritoneal cells from WT or renin KO mice. Peritoneal cells were plated in a 6 well dish in DMEM + 5% FBS for 60 minutes. The floating cells were removed, and the adherent cells were isolated using a cell scraper. Cells were then incubated with Salmonella at various cell : bacteria ratios (as indicated in the text) for 2 hours at 37°C. After 2 hours, cells were lysed with 0.02% SDS/PBS, and bacteria were transferred to Eppendorf tubes. Bacteria were then plated in agar dishes and grown overnight, and bacteria colony forming units were quantitated.

### Statistics

Statistical significance between groups was evaluated using the paired t test. Differences were considered statistically significant at *P<0.05 levels. Bar graphs are expressed as mean+/− s.e.m.

### Study Approval

All studies were performed in accordance with the National Institutes of Health Guide for the Care and Use of Laboratory Animals and were approved by the Animal Care and Use Committee of the University of Virginia.

## AUTHOR CONTRIBUTIONS

BCB, AES and TCM designed experiments, conducted experiments, acquired data, and analyzed data.WAV and VKN, conducted experiments, acquired data, and analyzed data. MLSSL and RAG designed and supervised research studies and wrote the manuscript.

## ACKNOWLEDGEMENTS

We thank Fang Xu and Tiffany Southard for expert technical assistance and animal care. We are grateful to the Child Health Research Center, the Flow Cytometry Core Facility and the Microscopy Core Facility at the University of Virginia, and Marieke Jones (Research Data Specialist at the Claude Moore Health Science Library) for technical assistance and expert advice. This research was supported the Karen Jargowsky Research Fund (BCB), and the National Institutes of Health (DK102914 to BCB, DK091330, DK096373, and DK116196 to MLSSL, and DK096373, DK116718, and HL066242 to RAG).

