## Supplemental Data for "A primitive type of renin-expressing lymphocyte protects the organism against infections"

### METHODS

**Flow Cytometry:** To characterize the immunophenotype of renin-lineage cells from reporter mice, flow cytometry analysis was performed. Single cell suspensions were obtained from the bone marrow, spleen, peripheral blood, and peritoneal fluid. Femurs were isolated and flushed with PBS+5% fetal bovine serum (FBS), and cells were passed through a 70 µm cell strainer. Spleens were passed through a 70 µm cell strainer using a syringe plunger to gently disrupt the tissues. Cells were rinsed through the strainer with PBS+FBS. Cell suspensions were treated with red blood cell lysis buffer, resuspended in PBS+FBS, counted using the Cellometer mini (Nexcelcom Bioscience, Lawrence, MA, USA) and distributed into microcentrifuge tubes for antibody labeling against surface markers (Supplemental Table 1). Antibodies were added at predetermined concentrations and incubated at room temperature for 20 minutes. Immunophenotyping was performed in the University of Virginia Flow Cytometry Core laboratory using a Fortessa cytometer, and data were analyzed with the FCSExpress program (De Novo Software, Los Angeles, CA, USA). For embryos, single cell suspensions were obtained from the yolk sac, fetal liver, and fetal spleen by passing tissues through a cell strainer as described above.

**RNA Extraction and Polymerase Chain Reaction Analysis:** Total RNA was isolated from the yolk sacs and livers of mouse embryos using Trizol extraction (Life Technologies, Grand Island, NY, USA) according to manufacturer's instructions. Complementary DNA (cDNA) was prepared from 2 µg RNA using Maloney murine leukemia virus reverse transcriptase (Life Technologies) and an oligo(dT) primer according to the manufacturer's instructions. PCR was performed on 2 µl cDNA using Taq DNA polymerase (Promega, Madison, WI, USA) in an Eppendorf thermocycler.

**Transplant studies:** Fetal liver cells from *Ren1*$^{dcre/+}$*;mTmG* mice were isolated (as above) and transplanted into adult WT mice after being treated with radiation (13 Gy divided into 2 fractions) via tail vein injection. Adult recipient mice were sacrificed 2 weeks later, tissues were harvested, and the percentage / identity of renin-lineage cells were determined by flow cytometry.

**Membrane Immuno Assay (MIA):** This was performed by culturing cells on a nitrocellulose membrane(1) followed by an immunoassay for renin modified based on the dot immunoassay protocols described previously(2). Briefly, cells were grown in low density on 0.22µ nitrocellulose membranes (Bio Rad) and cultured for 48h at 37ºC/5% $CO_2$ in six well plates. Cells were fixed on the membranes by baking them at 100ºC for 30 minutes followed by 2% PFA treatment for 30 minutes at room temperature (RT). The membranes were subsequently immune assayed for renin with all the steps performed with gentle shaking at RT. After fixing, membranes were washed with 0.1M Tris buffer; pH 7.4 + 0.05% tween (Buffer I) and further treated with 3% hydrogen peroxide for 15 minutes to quench the endogenous peroxide activity. Blocking was performed with 3% protease free bovine serum albumin (BSA) in 0.1M Tris buffer (pH7.4) for an hour. Primary antibody for renin (affinity purified rabbit polyclonal) at 1:2500 dilution in buffer I was added to the membrane and incubated overnight at RT. A biotinylated anti-rabbit secondary antibody kit **(**Vectasatin – Elite, ABC Reagent kit, Vector Laboratories) was used to detect and amplify the primary antibody signals, following the manufacturer's protocol. After treating with avidin-biotin solution, membranes were washed in buffer I and placed in a peroxidase-free chamber containing tetramethylbenzadine warmed to RT (TMB membrane peroxidase substrate solution, Kirkeguard and Perry Laboratories) and stored in dark.

**Immunofluorescence:** Immunofluorescence was performed on frozen sections. Imaging of $GFP^+$ cells from various tissues was done using a Leica DFC310 FX digital camera connected to a Leica DFC 480 microscope.

**SEM:** Peritoneal cells were isolated from reporter mice and cultured in a 6 well plate with 20 x 20 mm coverslips in the wells. Peritoneal cells were incubated for 1 hour, and then the supernatant was removed. 500 µl of 042 bacteria were added to each well, and the cells / bacteria were incubated for 2 hours. Plates were then centrifuged at 1400 RPM for 10 minutes, and the supernatant was removed. 4% paraformaldehyde was added to fix the cells / bacteria, and samples were stored at 4ºC overnight. Scanning electron microscopy was performed by the University of Virginia Microscopy Core Facility.

**Phagocytosis Assay:** Peritoneal cells were collected from $Ren1^{dcre/+}$*;mTmG* mice as described above. Cells were plated in a 6 well plate and incubated for 90 minutes. pHrodo E. coli BioParticles (Life Technologies, Eugene, OR, USA) were added to the wells and incubated for 2 hours. Cells were then collected from the wells, washed with PBS+FBS, centrifuged, and resuspended. Aliquots were labelled with antibodies (above) for flow cytometry analysis.

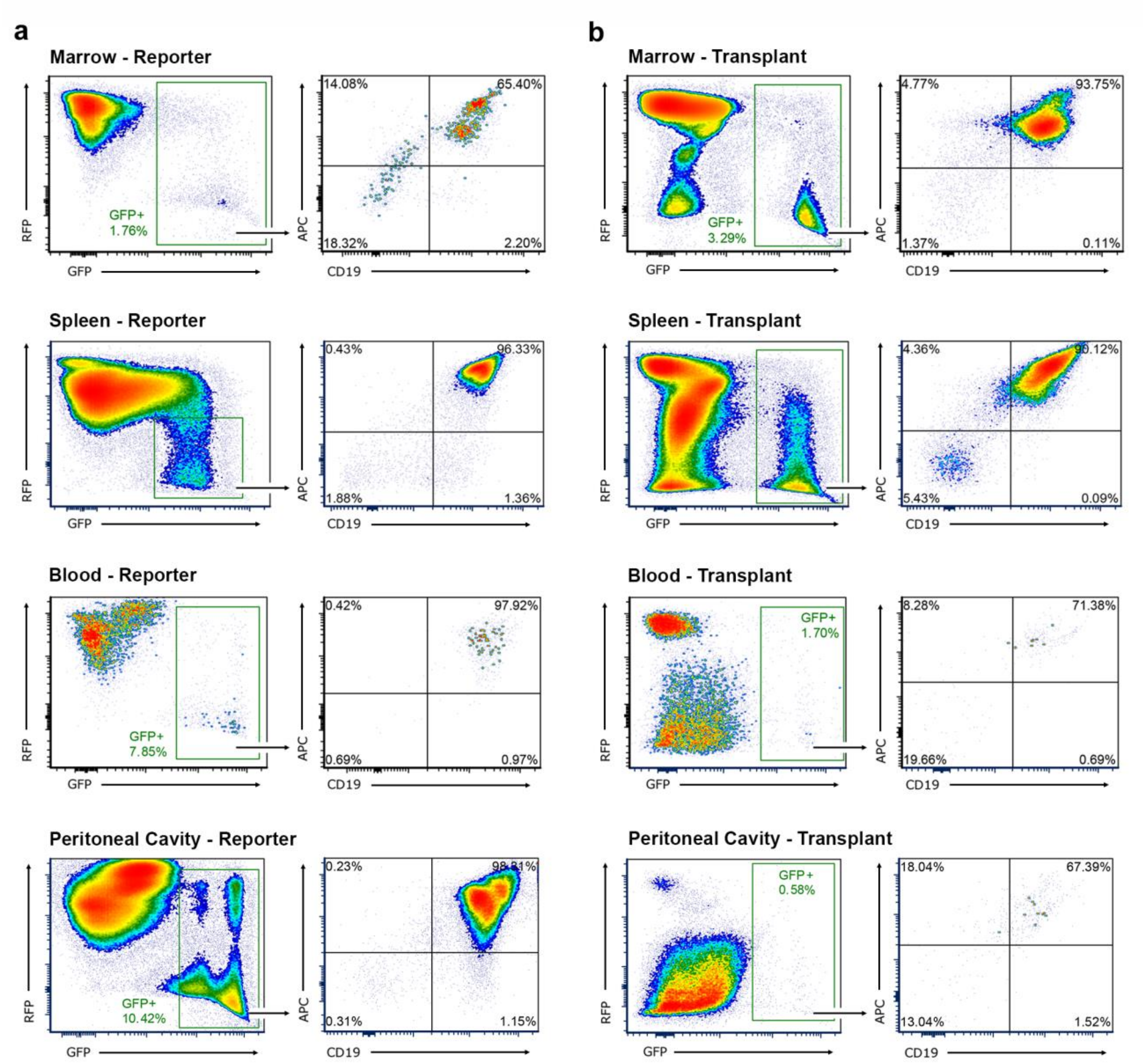


**Supplemental Figure 1. Transplant studies confirm the fate of renin-expressing progenitors during fetal life.**

a. Renin-lineage ($GFP^+$) cells were isolated from the bone marrow, spleen, blood, and peritoneal cavity of adult reporter mice (*Ren1*$^{dcre/+}$*; mTmG*). These cells were determined to be positive for the B cell surface markers CD19 and B220.

b. Fetal livers were isolated from E16.5 *Ren1*$^{dcre/+}$*;mTmG* embryos (where GFP marks renin-expressing cells and descendants). These cells were injected into the tail vein of irradiated adult wildtype hosts. Transplant recipients were studied 3 weeks after transplant, and engraftment was determined by the percent of GFP/RFP in the host organs. There was excellent engraftment in the bone marrow, but reduced engraftment in peripheral tissues. In all tissues, the transplanted renin lineage ($GFP^+$) cells mirrored renin lineage cells in adult reporter mice, expressing CD19 and B220.

**Supplemental Table 1. Antibodies used for flow cytometry**

| Antibody | | Fluorochrome | Concentration |
|---|---|---|---|
| CD19 | Pan B cell marker expressed on both B-1 and B-2 B cells | APC | 0.75 µg per $10^6$ cells |
| B220 | Pan B cell marker expressed highly on B-2 B cells and more dim on B-1 B cells | APC / Cy7 | 1 µg per $10^6$ cells |
| CD11b | Expressed on B-1 B cells | PerCP / Cy5.5 | 1 µg per $10^6$ cells |
| CD5 | Expressed on B-1a B cells but not B-1b B cells | Brilliant Violet 421 | 5 µl per $10^6$ cells |
| CD43 | Expressed on Pro-B cells and B-1 B progenitor cells | PE / Cy7 | 0.25 µg per $10^6$ cells |
| CD23 | Expressed on mature B-2 Bcells but not on B-1 B cells | PE / Cy7 | 0.25 µg per $10^6$ cells |
| Lineage Cocktail | Cocktail of antibodies against mature hematopoietic cells including B cells, T cells, Granulocytes, Monocytes, NK cells and Erythrocytes | Brilliant Violet 421 | 20 µl per $10^6$ cells |
